# LUZP1 regulates the constriction velocity of the contractile ring during cytokinesis

**DOI:** 10.1101/2022.07.04.498656

**Authors:** Toshinori Hyodo, Eri Asano-Inami, Satoko Ito, Mai Sugiyama, Akihiro Nawa, Md. Lutfur Rahman, Muhammad Nazmul Hasan, Yuko Mihara, Vu Quang Lam, Sivasundaram Karnan, Akinobu Ota, Shinobu Tsuzuki, Michinari Hamaguchi, Yoshitaka Hosokawa, Hiroyuki Konishi

## Abstract

There has been a great deal of research on cell division and its mechanisms; however, its processes have not yet been entirely elucidated. To find novel proteins that regulate cell division, we performed the screening using siRNAs and/or the expression plasmid of the target genes and identified leucine zipper protein 1 (LUZP1). Recent studies have shown that LUZP1 interacts with various proteins and stabilizes the actin cytoskeleton; however, the function of LUZP1 in mitosis is not known. In this study, we found that LUZP1 colocalized with the chromosomal passenger complex (CPC) at the centromere in metaphase and at the central spindle in anaphase and that these LUZP1 localizations were regulated by CPC activity and kinesin family member 20A (KIF20A). Mass spectrometry analysis identified that LUZP1 interacted with death-associated protein kinase 3 (DAPK3), one regulator of the cleavage furrow ingression in cytokinesis. In addition, we found that LUZP1 also interacted with myosin light chain 9 (MYL9), a substrate of DAPK3, and comprehensively inhibited MYL9 phosphorylation by DAPK3. In line with a known role for MYL9 in the actin-myosin contraction, LUZP1 suppression accelerated the constriction velocity at the division plane in our timelapse analysis. Our study indicates that LUZP1 is a novel regulator for cytokinesis that regulates the constriction velocity of the contractile ring.

## Introduction

The timing and location of protein activation and inactivation during cell division are highly regulated by post-translational modifications, such as protein phosphorylation and ubiquitination, to ensure the accurate progression of cell division (Lara-Gonzalez et al., 2021; Nasa et al., 2018; Vagnarelli, 2021). The chromosomal passenger complex (CPC) is an essential kinase complex for mitosis and is required for various mitotic events, such as chromosome condensation, kinetochore-microtubule attachment, spindle assembly checkpoint, cleavage furrow ingression, and abscission (Carmena et al., 2012; Krenn and Musacchio, 2015; McVey et al., 2021; Petsalaki and Zachos, 2021; Salaun et al., 2008). CPC is composed of four subunits: aurora kinase B (AURKB); inner centromere protein (INCENP); baculoviral IAP repeat-containing 5 (BIRC5; also known as Survivin); and cell division cycle associated 8 (CDCA8; also known as Borealin). AURKB is an enzymatic component of CPC and the remaining three are regulatory and targeting components that are essential for the proper kinase activity and localization of CPC. The key features of CPC are that its localization changes from the centromere to the central spindle when chromosomes segregate and that effective phosphorylation is only performed at its localizing area (Carmena et al., 2012).

Cytokinesis, the final step of cell division, begins in anaphase, with its cleavage furrow ingression regulated by CPC. After chromosome segregation, the bundles of an antiparallel microtubule, called the central spindle, are organized and the actin-myosin (actomyosin)-based contractile ring is assembled on the inner face of the plasma membrane at the division plane. Subsequently, the ring constricts when the actomyosin slides; thereby, each bundle of the central spindle is compacted to form the midbody on the intercellular bridge. Finally, the bridge is physically disconnected into two daughter cells (Husser et al., 2021; Leite et al., 2019). CPC on the central spindle promotes the recruitment and phosphorylation of the centralspindlin complex, thus stabilizing the central spindle and recruiting factors for cleavage furrow ingression (Carmena et al., 2012). The constriction of the contractile ring is mainly regulated by myosin II, which is composed of two myosin heavy chains (MHCs), two myosin essential light chains (ELCs; MLCs), and two myosin regulatory light chains (RLCs; MLCs) (Mishra et al., 2013; Sitbon et al., 2020; Yu et al., 2016). Furthermore, the motor activity of myosin II is regulated by the phosphorylation of RLCs by several kinases, such as myosin light chain kinases (MLCKs; MYLKs), Rho-associated coiled-coil containing protein kinases (ROCKs), and death-associated protein kinases (DAPKs) (Takeya et al., 2014; Yu et al., 2016). Myosin light chain 9 (MYL9), also known as LC20, is one RLC. Previous studies have reported that DAPK3, also known as ZIPK, directly phosphorylates MYL9 and regulates the actomyosin contraction in vascular smooth muscle via MYL9 phosphorylation (Deng et al., 2019; Moffat et al., 2011). In addition, recent studies have shown that DAPK3 also regulates cleavage furrow ingression in cytokinesis (Hamao et al., 2020; Hosoba et al., 2015; Ono et al., 2020).

Despite a great deal of research on cell division and its many mechanisms, its processes are not yet fully understood. To find novel proteins that regulate cell division, we screened and identified leucine zipper protein 1 (LUZP1), which contains three leucine zipper motifs and a coiled-coil domain at the N-terminus (Fig. 1A; Lee et al., 2001; Sun et al., 1996; Wang and Nakamura, 2019). The protein function of LUZP1 has long been unknown, but it is becoming rapidly apparent: LUZP1 mainly localizes to the actin filaments and stabilizes the actin cytoskeleton (Gonçalves, 2022; Wang and Nakamura, 2019); LUZP1 localizes at the centrosome and restricts primary cilia formation (Bozal-Basterra et al., 2020; Gonçalves et al., 2020); embryos of LUZP1 null knockout (KO) mice display cardiovascular and cranial defects with prenatal death (Hsu et al., 2008); LUZP1 genomic aberrations are detected in several types of cancer and syndromes (Bhat et al., 2022; Bozal-Basterra et al, 2021; Dong et al., 2021; Edwards et al., 2019; Poel et al., 2019; Zaveri et al., 2014); And LUZP1 interacts with microtubule and various other proteins and complexes (Fig. 1A; Krebs et al., 2010; Liu et al., 2017; Niri et al., 2021; Yano et al., 2020). Recent studies have also reported that LUZP1 localizes at the midbody during telophase (Bozal-Basterra et al., 2020; Gonçalves et al., 2020); however, no further information has been reported and the function of LUZP1 during mitosis is not yet known. In this study, we describe the novel function of LUZP1 during mitosis and present new insight for cell division research.

**Figure 1.**
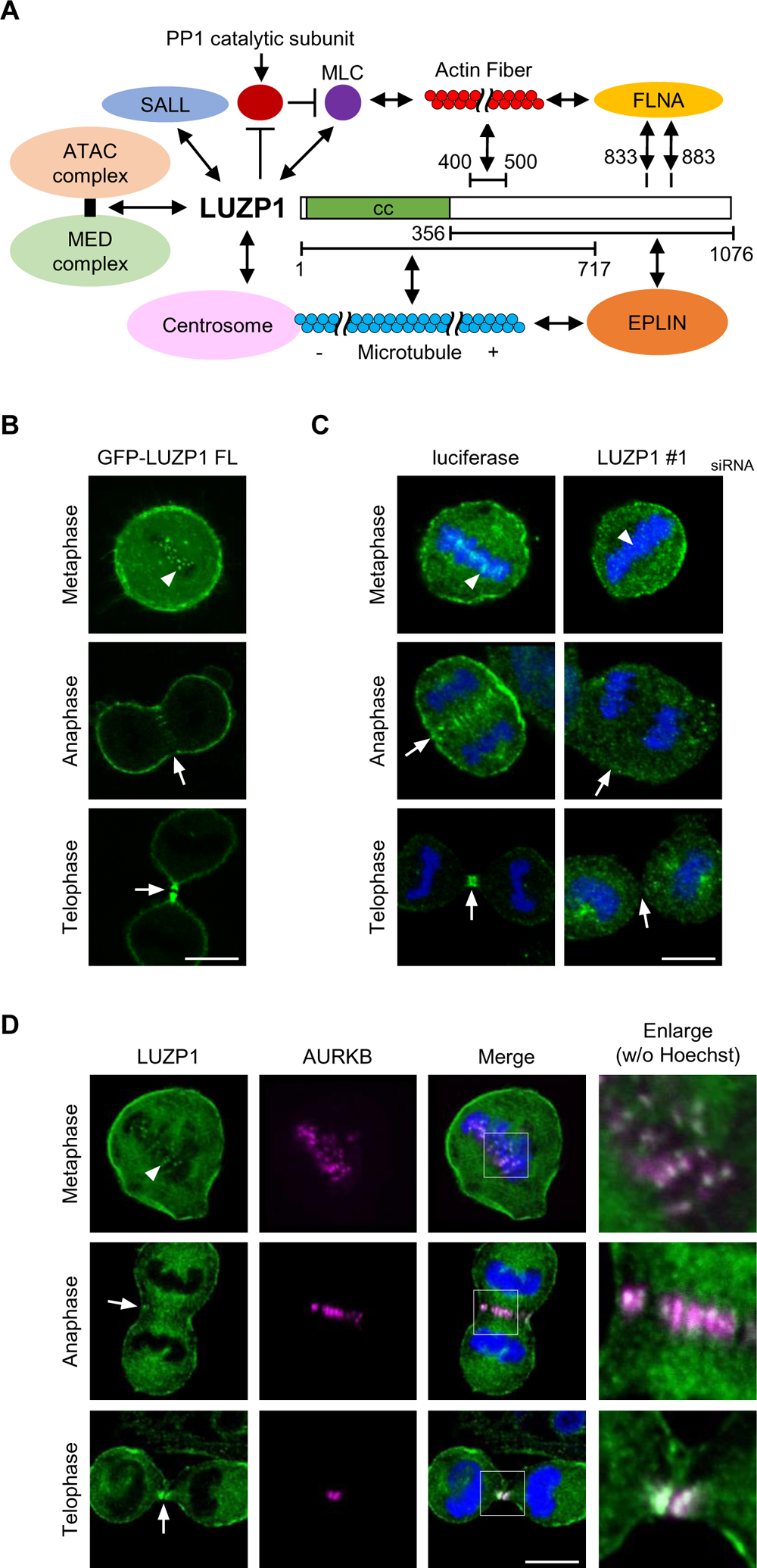
LUZP1 localizes at the inner centromere in metaphase, at the central spindle in anaphase, and at the midbody in telophase respectively. (A) Scheme: The two-way arrows indicate the proteins or complexes that interact with LUZP1; The bar range shows the interacting region of LUZP1. The abbreviations are as follows: PP1, Protein Phosphatase 1; MLC, myosin light chain; FLNA, filamin A; EPLIN (LIMA1), LIM domain and actin-binding 1; SALL, spalt-like transcription factor; ATAC, Ada-Two-A-containing; MED, Mediator; cc, coiled-coil domain. (B) HeLa cells were transfected with a plasmid encoding GFP-tagged full-length LUZP1 (GFP-LUZP1 FL), and the cells were observed alive. (C) HeLa cells were transfected with siRNAs against luciferase or LUZP1, and 72 h later the cells were immunostained with anti-LUZP1 antibodies and Hoechst. (D) HeLa cells were immunostained with anti-LUZP1 and anti-AURKB antibodies and Hoechst. (B) (C) (D) The arrowheads and the arrows indicate the centromere and the central spindle/midbody, respectively. The enlarged image shows the square area (scale bar, 10 μm).

## Results

### LUZP1 localizes at the inner centromere in metaphase, at the central spindle in anaphase, and at the midbody in telophase respectively

During mitosis, numerous proteins are phosphorylated by kinases, such as cyclin-dependent kinase 1 (CDK1) and CPC (Dephoure et al., 2008; Enserink and Kolodner, 2010; Kim et al., 2022; Salaun et al., 2008; Vagnarelli, 2021). Although the key kinases involved in phosphorylation have been well researched, the role of the phosphorylated proteins in mitosis remains unknown. Based on the findings by proteome analysis (Dephoure et al., 2008), we manually selected the candidates of mitosis-associated phosphorylated proteins, then performed the second screening using siRNAs and/or the expression plasmid of the target genes. The details of this screening and the mitotic functions of the other candidate proteins have been previously reported (Asano et al., 2013; Asano et al., 2014; Chen et al., 2015; Chen et al., 2017; Maeda et al., 2016; Hasegawa et al., 2013; Hyodo et al., 2012; Hyodo et al., 2016).

LUZP1 is a novel mitosis-related protein candidate found through our screening. Recent studies have shown that LUZP1 interacts with various proteins in interphase (Fig. 1A); however, the function of LUZP1 in mitosis is not yet known. To gain insight into the role of LUZP1 during mitosis, we examined its subcellular localization by transfecting a plasmid encoding, GFP-tagged, full-length LUZP1 (GFP-LUZP1 FL) into HeLa cells. Each phase of cell division was obtained by the release from the nocodazole-mediated prometaphase arrest (nocodazole-arrested and released). As shown in Fig. 1B, the live cell imaging showed GFP-LUZP1 FL at the centromere (arrowhead) during metaphase, at the central spindle/midbody (arrow) during anaphase and telophase, and at the cortical actin cytoskeleton under the plasma membrane throughout mitosis. In addition, GFP-LUZP1 FL was also observed at the centrosome during mitosis (Fig. 1B). To validate these findings, we generated anti-LUZP1 antibodies capable of detecting endogenous LUZP1. Hela cells were transfected with siRNAs against luciferase or LUZP1, nocodazole-arrested and released, and then immunostained with anti-LUZP1 antibodies. In luciferase siRNA-transfected cells, we observed endogenous LUZP1 at the same location as GFP-LUZP1 FL (Fig. 1C); whereas in the LUZP1 siRNA-transfected cells, these localizations disappeared (Fig. 1C). We also examined whether LUZP1 colocalizes with AURKB, since AURKB is known to localize at the inner centromere during metaphase and at the central spindle/midbody during anaphase and telophase. We found that endogenous LUZP1 colocalized with AURKB in each phase of cell division (Fig. 1D). These results indicate that LUZP1 localizes at the inner centromere during metaphase and at the central spindle/midbody during anaphase and telophase.

### The centromere and the central spindle localization of LUZP1 require CPC kinase activity and KIF20A, respectively

When chromosomes segregate, LUZP1 disappeared from the centromere and appeared at the central spindle (Fig. 1B-D). Interestingly, these LUZP1 localizations seem to translocate similar to CPC (Carmena et al., 2012; Krenn and Musacchio, 2015; McVey et al., 2021; Petsalaki and Zachos, 2021). Previous studies have demonstrated that CPC translocation from the centromere to the central spindle requires kinesin family member 20A (KIF20A), which is also known as MKLP2 (Adriaans et al., 2020; Kitagawa et al., 2013). To investigate the relationship between LUZP1 and CPC, GFP-LUZP1 FL plasmid was transfected into HeLa cells with siRNAs against luciferase or KIF20A. We observed GFP-LUZP1 FL at the centromere in both the luciferase and KIF20A siRNA-transfected cells during metaphase (Fig. 2A; arrowhead); whereas, the central spindle/midbody localization of GFP-LUZP1 FL disappeared during anaphase and telophase in the KIF20A siRNA-transfected cells only (Fig. 2A; arrow). This result indicates that the central spindle localization of LUZP1 requires KIF20A, which is similar to CPC. Since previous studies have reported that CPC also regulates the localization of some proteins and complexes (Carmena et al., 2012), we treated cells with VX-680, an aurora kinase-specific inhibitor, to examine the resulting LUZP1 localizations. Briefly, we transiently expressed DsRed-tagged full-length LUZP1 (DsRed-LUZP1 FL) and GFP-tagged full length AURKB (GFP-AURKB) together into HeLa cells, which were then treated with nocodazole. Afterwards, the cells were additionally treated with VX-680. Immediately after VX-680 treatment, DsRed-LUZP1 FL colocalized with GFP-AURKB at the centromere (Fig. 2B; arrowhead); however, DsRed-LUZP1 FL disappeared from the centromere 30 min after VX-680 treatment, whereas GFP-AURKB remained at the centromere (Fig. 2B; arrowhead). Furthermore, the cortical actin cytoskeleton localization of DsRed-LUZP1 FL was maintained at this time point (Fig. 2B). This result indicates that the centromere localization of LUZP1 requires CPC kinase activity. In addition, these results suggest that LUZP1 translocates from the centromere to the central spindle under the regulation of CPC and KIF20A.

**Figure 2.**
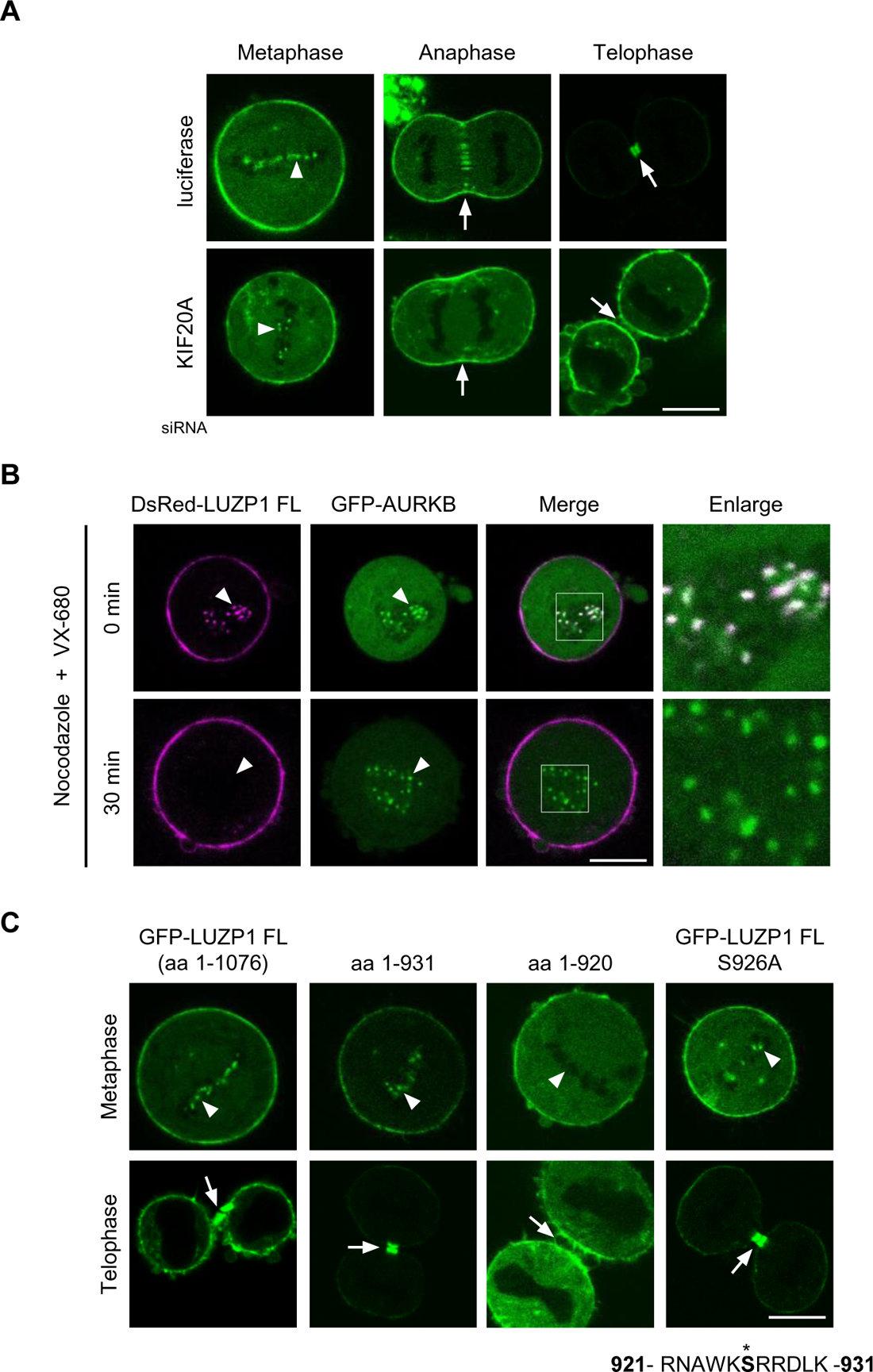
The centromere and the central spindle localization of LUZP1 require CPC kinase activity and KIF20A, respectively. (A) A plasmid encoding GFP-tagged full-length LUZP1 (GFP-LUZP1 FL) was transfected into HeLa cells together with siRNAs against luciferase or KIF20A, with live cell images taken 24 h later. (B) DsRed-LUZP1 FL and GFP-AURKB were transiently expressed into HeLa cells together, then the cells were treated with nocodazole for 14 h. Next, the cells were additionally treated with VX-680, an aurora kinase-specific inhibitor, with the live cells observed 0 or 30 min later. (C) HeLa cells were transfected with the indicated GFP-LUZP1 plasmid, with live cell images taken 24 h later. Character string and asterisk (*) show the amino acid (aa) sequence of LUZP1 and the phosphorylation site candidate, respectively. (A) (B) (C) The arrowheads and the arrows indicate the centromere and the central spindle/midbody, respectively. The enlarged image shows the square area (scale bar, 10 μm).

We next examined which LUZP1 region is required for the centromere and the central spindle/midbody localizations by transfecting GFP-LUZP1 FL or deletion mutants of LUZP1 into HeLa cells. As shown in Fig. 2C, the amino acid (aa) 1-931 LUZP1 was observed at the centromere and the midbody like GFP-LUZP1 FL, but the aa 1-920 LUZP1 was not observed at either location. This result indicates that only the 11 aa from 921 to 931 are critical for the centromere and the central spindle/midbody localizations. Among these 11 aa, Ser926 was the only phosphorylation site (Fig. 2C; character string). By generating the GFP-LUZP1 FL S926A plasmid, which substitutes Ser926 for alanine, and transfecting it into HeLa cells, we found that GFP-LUZP1 FL S926A still localized at the centromere and the midbody (Fig. 2C). We additionally generated other mutants which substituted the predicted phosphorylation site(s) in mitosis for alanine: GFP-LUZP1 FL S573A; S574A; T679A; S840A; S878A; and S891A; however, these mutants localized at both locations (Supplemental Fig. S1). These results indicate that the LUZP1 centromere localization does not require the phosphorylation of the aa 921-931 LUZP1, although it does require CPC kinase activity and the aa 921-931 LUZP1 (Fig. 2B and 2C). Although the above findings suggest that LUZP1 associates with CPC, we did not obtain any experimental results showing the interaction between LUZP1 and CPC (see discussion).

### LUZP1 overexpression induces binucleated cells

The protein expression changes essential for mitosis are known to induce mitotic failure, such as mitotic arrest, lagging chromosome, and multinucleated cells; however, recent studies have shown that LUZP1 suppression does not induce multinucleated cells (Bozal-Basterra et al., 2020; Gonçalves et al., 2020). To confirm that LUZP1 has a function in mitosis, HeLa cells were transfected with GFP-LUZP1 FL plasmid or LUZP1 siRNA. We found that binucleated (multinucleated) cells were slightly increased in GFP-LUZP1 FL overexpressed cells, (Fig. 3A and 3B). In contrast, binucleated cells were not increased in the LUZP1 siRNA-transfected cells, as previously reported (Fig. 3B; Supplemental Fig. S2A and S2B; Bozal-Basterra et al., 2020; Gonçalves et al., 2020). Since complications during cytokinesis are known to induce binucleated cells, these results suggest that LUZP1 relates to cytokinesis during mitosis, even if its function is not essential.

**Figure 3.**
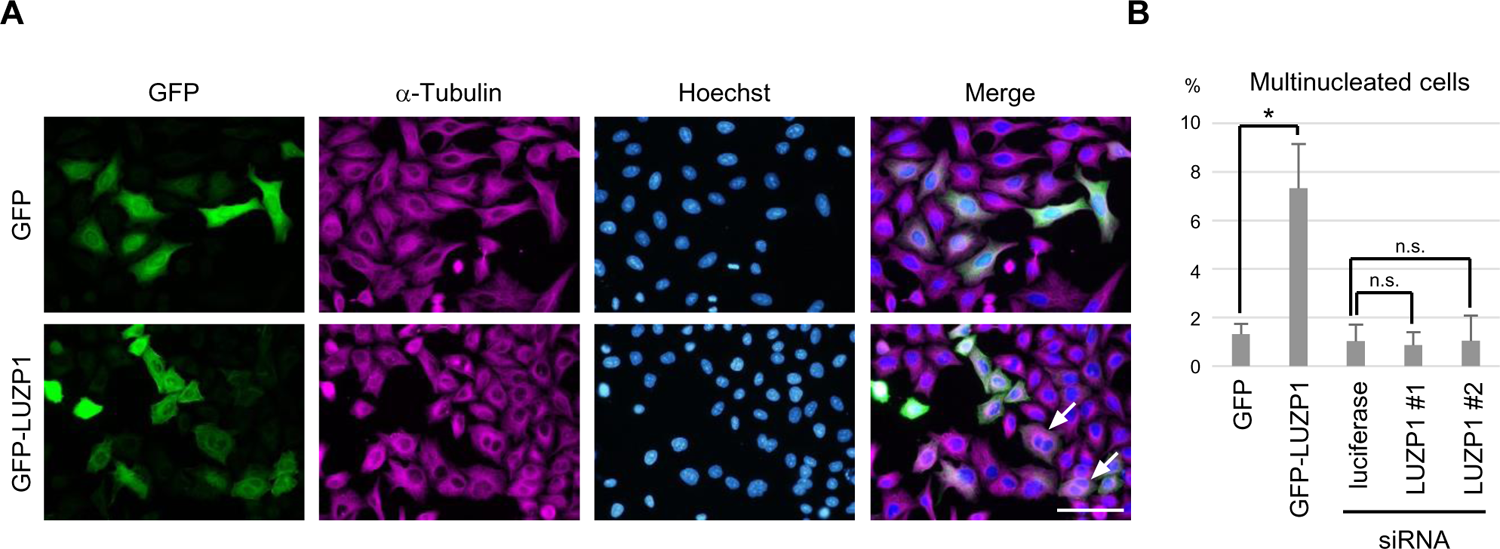
LUZP1 overexpression induces binucleated cells. (A) A plasmid encoding GFP-tagged full-length LUZP1 (GFP-LUZP1 FL) or GFP tag was transfected into HeLa cells. After 72 h, the cells were immunostained with anti-α-Tubulin and anti-GFP antibodies and Hoechst. The arrows indicate binucleated (multinucleated) cells (scale bar, 50 μm). (B) The ratio of multinucleated cells in (A) and (Supplemental Fig. S2A) was evaluated. Mean and SEM value of three independent experiments are shown, and more than 100 cells were evaluated for each experiment; *, *P* < 0.05; n.s., not significant.

### LUZP1 interacts with DAPK3

Recent studies have shown that LUZP1 interacts with various proteins during interphase (Fig. 1A), but these proteins during mitosis have not yet been reported. Thus, we used mass spectrometry to gain further insight into the function of LUZP1 during mitosis by producing 293T cells that constitutively expressed Flag-LUZP1 FL. The cells were then either nocodazole-arrested and released or untreated with nocodazole and immunoprecipitated with anti-Flag antibodies to identify the coprecipitated proteins. We detected DAPK3 in both cell samples (Fig. 4A; Supplemental Table). Although we were unable to determine the interacting protein in mitosis only, this result suggests that DAPK3 interacts with LUZP1 throughout the cell cycle. To confirm the interaction between LUZP1 and DAPK3 during mitosis, a plasmid encoding GFP-tagged full-length DAPK3 (GFP-DAPK3) was transfected into 293T cells together with a plasmid encoding Flag-tagged full-length or deletion mutants of LUZP1. Next, the cells were nocodazole-arrested and released, then immunoprecipitated with anti-Flag antibodies. As shown in Fig. 4B, GFP-DAPK3 coprecipitated with Flag-LUZP1 FL but not with Flag tag alone. In addition, GFP-DAPK3 coprecipitated with a coiled-coil (cc) domain of LUZP1 only (Flag-LUZP1 cc) and not with the cc domain deletion LUZP1 (Flag-LUZP1Δcc; Fig. 4B). These results indicate that the cc domain of LUZP1 was responsible for its interaction with DAPK3.

**Figure 4.**
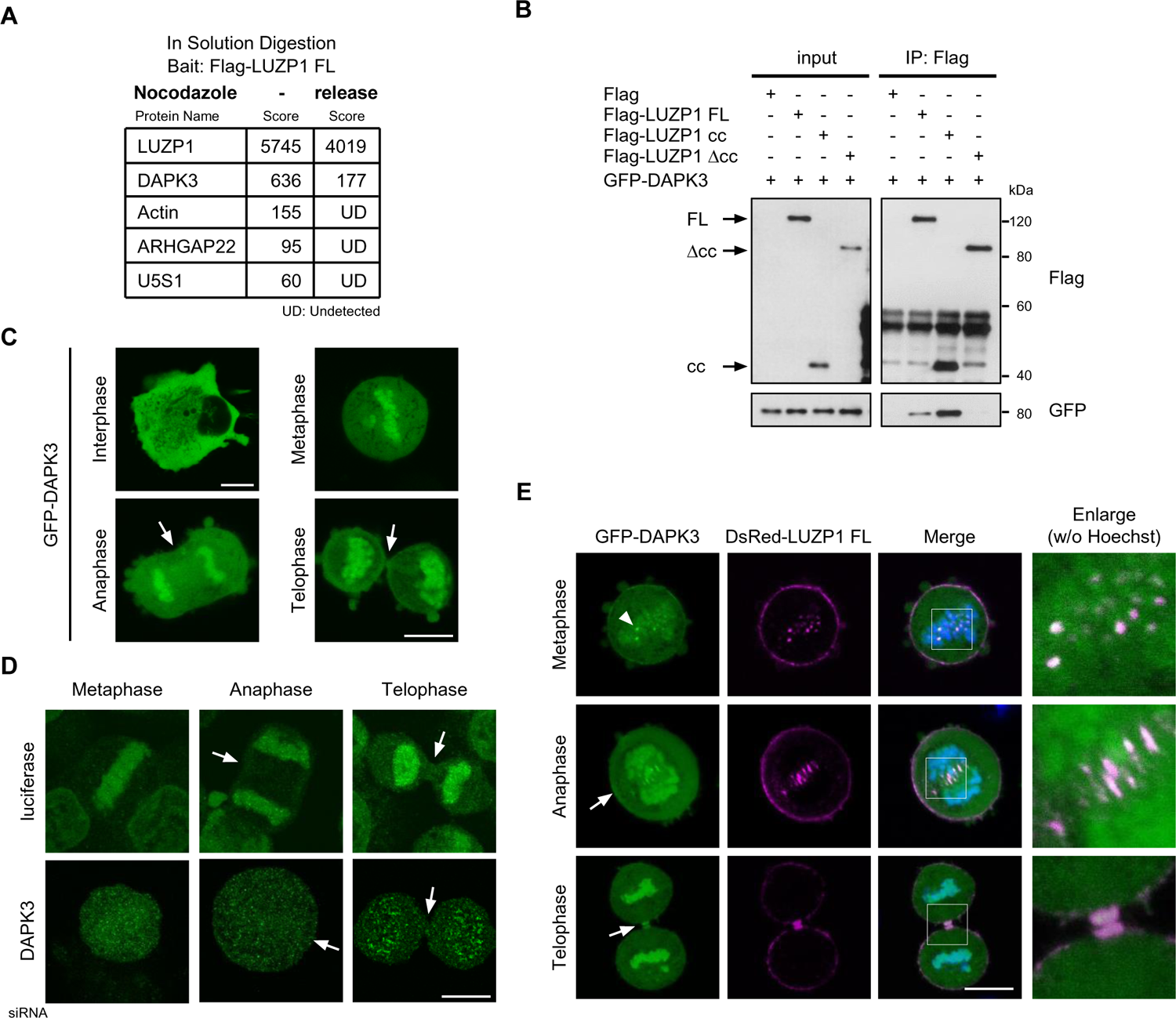
LUZP1 interacts with DAPK3. (A) A list of proteins identified by mass spectrometry analysis. (B) GFP-tagged full-length DAPK3 (GFP-DAPK3) and the indicated Flag-tagged full-length or deletion mutants of LUZP1 were transiently expressed into 293T cells together, then immunoprecipitated with anti-Flag antibodies. Next, the immunoprecipitants were immunoblotted with anti-Flag and anti-GFP antibodies. (C) GFP-DAPK3 was transiently expressed into HeLa cells, with live images taken 24 h later. (D) HeLa cells were transfected with siRNAs against luciferase or DAPK3, and the cells were immunostained with anti-DAPK3 antibodies 72 h later. (E) GFP-DAPK3 and DsRed-tagged full-length LUZP1 (DsRed-LUZP1 FL) were transiently expressed into HeLa cells together, with live images taken 24 h later. (C) (D) (E) The arrowheads and the arrows indicate the centromere and the central spindle/midbody, respectively. The enlarged image shows the square area (scale bar, 10 μm).

The above results suggest that LUZP1 interacts with DAPK3 in mitosis. To further investigate this relationship, we examined the subcellular localization of DAPK3 during mitosis by transiently expressing GFP-DAPK3 into HeLa cells, then performing nocodazole-arrest and release. We observed GFP-DAPK3 at the chromosome throughout mitosis, but not at the nucleus during interphase (Fig. 4C). To determine the localization of endogenous DAPK3 in mitosis, HeLa cells were transfected with siRNAs against luciferase or DAPK3 and immunostained with anti-DAPK3 antibodies. Endogenous DAPK3 was observed at the chromosome in the luciferase siRNA-transfected cells but not in the DAPK3 siRNA-transfected cells (Fig. 4D). These results indicate that DAPK3 certainly localizes at the chromosome during mitosis. Although the centromere is on the chromosome, this result indicates that the localization of DAPK3 only partially matches that of LUZP1. To confirm the association between DAPK3 and LUZP1 in the cells, GFP-DAPK3 and DsRed-LUZP1 FL were transiently expressed into HeLa cells together, which resulted in GFP-DAPK3 colocalizing with DsRed-LUZP1 FL at the centromere during metaphase and at the central spindle/midbody during anaphase and telophase in addition to the chromosome localization (Fig. 4E and Supplemental Fig. S3). Although the localization between LUZP1 and DAPK3 only partially match, the above results suggest that DAPK3 can interact with LUZP1 in a cell.

### LUZP1 regulates MYL9 phosphorylation by DAPK3

Previous studies have reported that DAPK3 regulates cleavage furrow ingression in cytokinesis and that DAPK3 directly phosphorylates MYL9, which regulates the actomyosin contraction (Takeya et al., 2014; Moffat et al., 2011; Deng et al., 2019). Interestingly, a recent study reported that LUZP1 interacts with one MLC (Yano et al., 2020). To confirm the relationships between LUZP1, DAPK3, and MYL9, we tested whether LUZP1 interacts with MYL9. Briefly, the GST-tagged full-length MYL9 (GST-MYL9) or GST tag alone proteins were produced in *E. coli*, then bound to glutathione agarose beads and purified (Fig. 5A). Flag-LUZP1 FL or Δcc was transiently expressed into 293T cells, then its extracts were mixed with the beads bound to GST-MYL9 or GST. As shown in Fig. 5A, both LUZP1 FL and Δcc were pulled down by GST-MYL9 but not by GST tag alone. This result indicates that LUZP1 also interacts with MYL9 at a different interaction region from DAPK3 (Fig. 5B). To validate whether LUZP1 regulates MYL9 phosphorylation by DAPK3, we next performed an *in vitro* kinase assay. Flag-LUZP1 FL or Δcc was transiently expressed into 293T cells, then bound to beads conjugated with anti-Flag antibodies and purified. The purified GST-MYL9 was eluted using reduced glutathione. Recombinant active DAPK3, GST-MYL9, and [γ-^32^P] labeled ATP were mixed with different amounts of Flag-LUZP1 FL or Δcc and incubated for 30 min. Next, the reaction mixtures were separated by SDS-PAGE gel and subjected to autoradiography. DAPK3 directly phosphorylated GST-MYL9, but not GST (Fig. 5C). We also found that MYL9 phosphorylation by DAPK3 was dose-dependently inhibited in both Flag-LUZP1 FL and Δcc (Fig. 5D and 5E). Even more interestingly, LUZP1 Δcc suppressed MYL9 phosphorylation more strongly than LUZP1 FL (Fig. 5D and 5E), although the same LUZP1 protein amount was added (Fig. 5F). To further investigate the differences between LUZP1 FL and Δcc, we measured the efficiency of the MYL9 phosphorylation by DAPK3. Recombinant active DAPK3, GST-MYL9, and [γ-^32^P] labeled ATP were mixed with a fixed amount of Flag-LUZP1 FL or Δcc, then the mixtures were incubated at 0, 10, 20, and 30 min, respectively. As shown in Fig. 5G and 5H, the velocity of MYL9 phosphorylation containing LUZP1 FL was faster than those containing LUZP1 Δcc. These results indicate that the interaction between LUZP1 and MYL9 prevents MYL9 phosphorylation by DAPK3; however, the interaction between LUZP1 and DAPK3 induces MYL9 phosphorylation in reverse.

**Figure 5.**
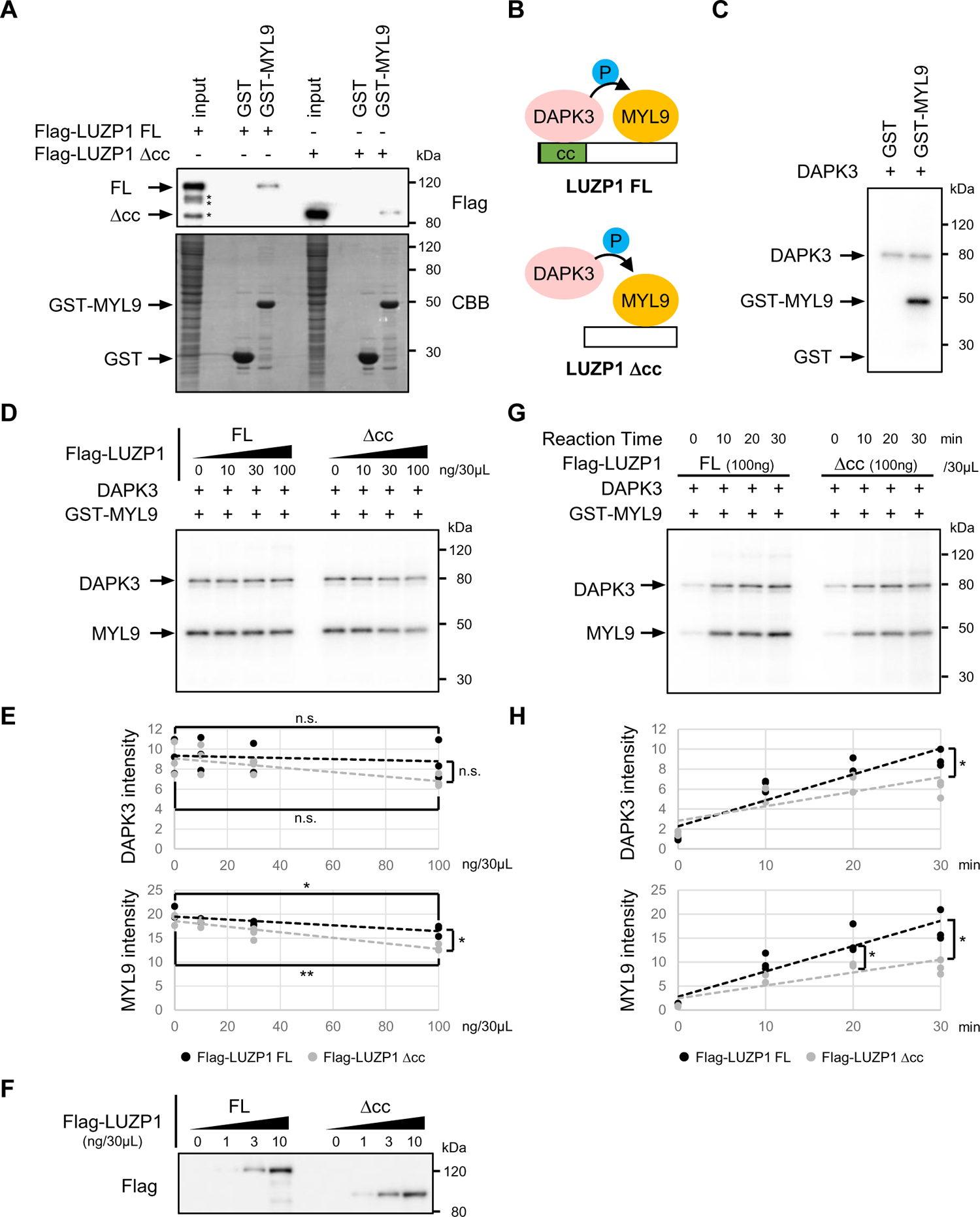
LUZP1 regulates MYL9 phosphorylation by DAPK3. (A) The purified GST-tagged full length MYL9 (GST-MYL9) and GST tag proteins were separated by an SDS-PAGE gel and stained with Coomassie Brilliant Blue (CBB). Flag-tagged full-length LUZP1 (Flag-LUZP FL) or the cc domain deletion LUZP1 (Flag-LUZP1 Δcc) was transiently expressed into 293T cells and its extracts were mixed with the beads bound to GST-MYL9 or GST, then the proteins were pulled down. The samples were immunoblotted with anti-Flag antibodies. The asterisks (*) indicate degraded Flag-LUZP1 FL proteins. (B) Scheme shows the interacting regions among LUZP1, DAPK3, and MYL9. (C) (D) (G) Active DAPK3 (10 ng), purified GST-MYL9 or GST (1 μg), and [γ^32^-P] ATP (1.0 μCi, 0.33 pmol) were mixed with the indicated amount of LUZP1 FL or Δcc, then incubated for 30 min or at the indicated time at 30°C. The reaction mixtures were separated by SDS-PAGE gel and subjected to autoradiography. (E) (H) Each dot shows the relative intensity of phosphorylated MYL9 or auto-phosphorylated DAPK3 in three independent kinase assays. Mean and SEM value of three independent experiments are shown; **, *P* < 0.01; *, *P* < 0.05; n.s., not significant. (F) One-tenth amount of Flag-LUZP1 FL and Δcc in (D) were immunoblotted with anti-Flag antibodies.

### LUZP1 regulates the constriction velocity of the contractile ring in cytokinesis

The above findings suggest that LUZP1 regulates the actomyosin contraction in cytokinesis by controlling the MYL9 phosphorylation by DAPK3. To confirm this hypothesis, we measured the constriction velocity of the contractile ring in cytokinesis via timelapse analysis. HeLa cells were transfected with LUZP1 siRNA or GFP-LUZP1 FL plasmid and the cells were nocodazole-arrested and released. Next, the timelapse imaging during cell division were taken for 3 sec intervals using confocal microscopy with an incubator. During this experiment, the GFP image was taken only at the initial point since the short interval laser exposure for GFP excitation was harmful to the dividing cells. In addition, we measured the time required for the division plane to be halved in diameter starting immediately after chromosome segregation to perform the statistical analysis, because the time required for the abscission, the final step of cytokinesis, is very unstable. We found that LUZP1 siRNA-transfected cells had an accelerated constriction velocity of the contractile ring compared to the luciferase siRNA-transfected cells (Fig. 6A, 6B, 6C, and Supplemental Movie S1-S3). In contrast, the velocity was delayed in the GFP-LUZP1 FL transfected cells compared to the GFP tag transfected cells (Fig. 6A, 6B, 6C, and Supplemental Movie S4, S5). These results indicate that LUZP1 suppresses the constriction velocity of the contractile ring during cytokinesis.

**Figure 6.**
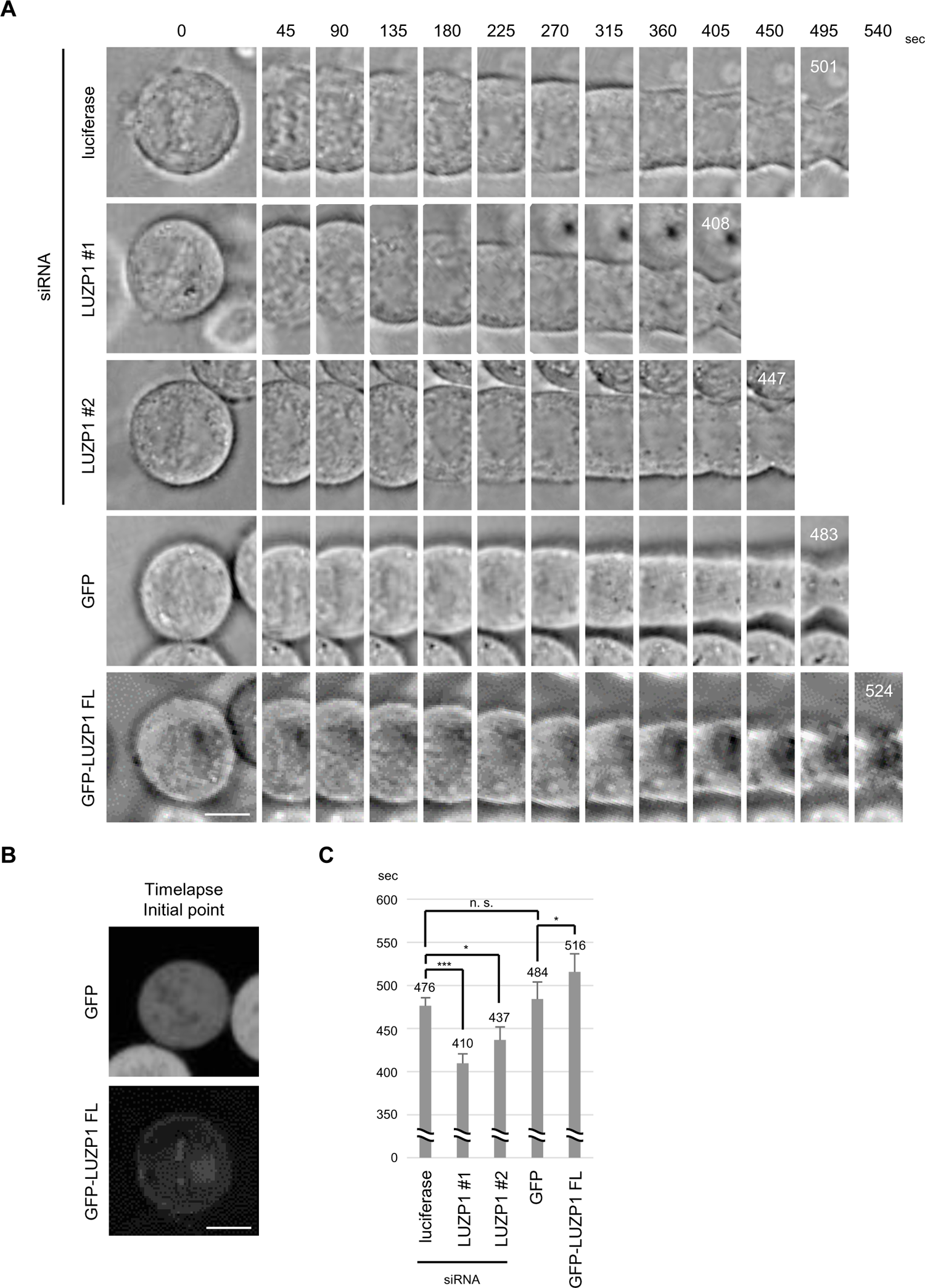
LUZP1 regulates the constriction velocity of the contractile ring in cytokinesis. (A) HeLa cells were transfected with the indicated siRNA or plasmid. Timelapse images were taken for 3 sec intervals using confocal microscopy with an incubator. The images were arranged every 45 sec. The last white number indicates the time at which the contractile ring diameter was halved (scale bar, 10 μm). (B) The GFP or GFP-tagged full-length LUZP1 (GFP-LUZP1 FL) fluorescence was taken only at the initial point to pick up the GFP expressing cells in (A). (C) The graph shows the time required for the contractile ring to be halved in diameter starting immediately after chromosome segregation. Mean and SD value are shown; ***, *P* < 0.001; *, *P* < 0.05; n.s., not significant. Measured cell numbers were as follows: luciferase, n = 32; LUZP1 #1, n = 39; LUZP1 #2, n = 18; GFP, n = 10; GFP-LUZP1 FL, n = 11.

## Discussion

In this study, we show that LUZP1 localizes at the inner centromere and the central spindle during metaphase and anaphase, respectively (Fig. 1B-D). In addition, we also provide evidence that LUZP1 probably translocates from the centromere to the central spindle under CPC and KIF20A regulation (Fig. 2A and 2B). These results suggest that LUZP1 is a new component of CPC similar to regulator of chromosome condensation 2 (RCC2; also known as TD-60) (Mollinari et al., 2003; Papini et al., 2015). To confirm the interaction between LUZP1 and CPC, we performed a coimmunoprecipitation analysis using a plasmid encoding Flag-tagged full length each component of CPC (Flag-AURKB, Flag-INCENP, Flag-BIRC5, or Flag-CDCA8). We found that GFP-LUZP1 FL clearly coprecipitated with Flag-AURKB and faintly with Flag-INCENP and Flag-BIRC5 but not with Flag-CDCA8 and Flag tag (Supplemental Fig. S4A). It remains unclear which CPC component primarily interacts with LUZP1, because the protein expression levels could not be adjusted due to differences in overexpressed protein cytotoxicity. This result suggests that LUZP1 associates with CPC during mitosis. To further confirm this interaction, we examined whether LUZP1 dissociates with CPC when its centromere localization is prevented and found that GFP-LUZP1 FL coprecipitated with Flag-AURKB, even when VX-680 prevented LUZP1 centromere localization (Supplemental Fig. S4B). Furthermore, GFP-AURKB also coprecipitated with Flag-aa 1-920 LUZP1, the centromere unlocalized mutant, as it did with Flag-LUZP1 FL and Flag-aa 1-931 LUZP1 (Supplemental Fig. S4C). These results contradict the finding that aa 1-920 LUZP1 disappeared from the centromere, even though it still interacted with AURKB. To solve this puzzle, we tried to reveal which CPC component directly interacts with LUZP1, but we failed to produce the recombinant LUZP1 protein via *E. coli* or the baculovirus expression system, as previously reported (Wang and Nakamura, 2019). The inconsistencies between the interactions and localizations of LUZP1 have also been observed for other proteins, such as DAPK3 and EPLIN (Fig. 4B-E; Gonçalves et al., 2020; Gonçalves, 2022). The reason for this contradiction is still unknown, but it is possible that the affinity for binding between LUZP1 and its interactant is greatly reduced when LUZP1 is localized at the structures compared to when LUZP1 is unlocalized in the cytosol or in cell extracts. Our findings clearly show that LUZP1 localizes at the centromere and the central spindle/midbody during mitosis (Fig. 1B-D); however, further analysis is needed to determine whether LUZP1 is a novel component of CPC.

Our kinase assay indicated that LUZP1 inhibits the phosphorylation of MYL9 by DAPK3 (Fig. 5D-F); however, a recent study reported that LUZP1 promotes the MLC phosphorylation state by preventing the function of protein phosphatase 1 (Yano et al., 2020). Although these results seem contradictory, the function of LUZP1 likely easily altered depending on the interactant involved, since LUZP1 does not have a functional domain and interacts with various proteins. In fact, our kinase assay indicates that the interaction between LUZP1 and MYL9 prevents the phosphorylation of MYL9 by DAPK3, but the interaction between LUZP1 and DAPK3 induces MYL9 phosphorylation in reverse (Fig. 5D-H). These findings suggest that one function of LUZP1 is to support protein-protein reactions at its localizing area as a scaffold protein.

In this study, we detected that LUZP1 suppresses the constriction velocity of the contractile ring during cytokinesis (Fig. 6A-C). Even when the constriction velocity was accelerated by LUZP1 suppression, cell division progressed and was completed normally (Fig. 3A and 3B; Supplemental Fig. S2A). It remains unknown why cells have the unnecessary delay in contraction during cytokinesis via LUZP1. Recent studies have reported that LUZP1 expression is downregulated in several types of cancer and that LUZP1 can suppress the proliferation of some cancer cells (Dong et al., 2021; Bozal-Basterra et al, 2021). These findings suggest that LUZP1 may contribute to genome stability by preventing a failed chromosome distribution by delaying cytokinesis.

Here, we described the novel function of LUZP1 during mitosis in mammalian cells. We showed that LUZP1 colocalizes with CPC at the inner centromere in metaphase, at the central spindle in anaphase, and at the midbody in telophase respectively, and that these localizations require CPC kinase activity and KIF20A. In addition, we provided evidence that LUZP1 interacts with both DAPK3 and its substrate, MYL9 and that LUZP1 regulates the phosphorylation of MYL9 by DAPK3 depending on the interaction of these proteins. Furthermore, we showed that LUZP1 regulates the constriction velocity of the contractile ring during cytokinesis. We believe that the novel function of LUZP1 during mitosis contributes to the advancement of cell division research.

## Acknowledgments

We would like to thank the staff at the Institute of Comprehensive Medical Research, Division of Advanced Research Promotion and Division of Radioisotope Research at Aichi Medical University, and Kentaro Taki and colleagues at the Division for Medical Research Engineering, Nagoya University Graduate School of Medicine, for providing technical assistance. This work was partially supported by Grants-in-Aid for Scientific Research (KAKENHI) from the Japan Society for the Promotion of Science (JSPS; 18K14703 to T.H., 19K09292 and 22K08985 to S.K., 21K08426 to A.O., 22H02856 to S.T., 19K08668 and 22K08294 to Y.H., and 20K06613 to H.K.), The Nitto Foundation (to T.H.), Takeda Science Foundation (to T.H.), and The Hori Science and Arts Foundation (to T.H. and A.O.). M.L.R. and M.N.H. are supported by the Japanese Government (MEXT) Scholarship for Research Students.

## Author contributions

T.H. conceived and designed the project. T.H. performed most of the experiments and analyzed data in this study. E.A-I. performed the partial experiments. S.I. guided T.H. in the experiment of mass spectrometry. M.S. and A.N. supported the analysis partially. M.L.R., M.N.H., Y.M., V.Q.L., S.K., A.O., S.T., M.H., Y.H., and H.K. provided insights into the project. T.H. wrote the manuscript with a significant contribution from H.K.

## Materials & methods

### Cells, antibodies, and chemicals

HeLa (RCB3680) and 293T (RCB2202) cells were obtained from RIKEN Bioresource Center (Ibaraki, Japan). Both cells were propagated in Dulbecco’s Modified Eagle’s Medium (D-MEM; Fujifilm, Osaka, Japan) supplemented with 10 % Fetal Bovine Serum (FBS; Nichirei Bioscience, Tokyo, Japan) and 1 % Penicillin-Streptomycin (Fujifilm).

Anti-LUZP1 antibodies were generated by injecting 200 μg of GST-LUZP1 (aa 262-427) mixed with Freund adjuvant (Merck, Darmstadt, Germany) into a rabbit every two weeks totally four times. After serum was obtained, the anti-LUZP1 antibodies were purified using an NHS-activated column (Cytiva, Marlborough, MA, USA) coupled with aa 262-427 GST-LUZP1. Anti-GST antibodies were eliminated using recombinant GST. Other antibodies and chemicals were obtained from the following manufacturers: anti-AIM-1 (AURKB) mouse monoclonal (6/AIM-1) antibodies (#611082) (BD Biosciences, Franklin Lakes, NJ, USA); anti-GFP polyclonal antibodies (#598) (Medical & Biological Laboratories, Tokyo, Japan); anti-α-Tubulin mouse monoclonal (DM1A) antibodies (#T6199) (Merck); anti-DAPK3 polyclonal antibodies (#PA5-90193) (Thermo Fisher Scientific, Waltham, MA, USA); anti-FLAG mouse monoclonal (M2) antibodies (#F3165) (Merck); Hoechst 33342 solution (Dojindo, Kumamoto, Japan); Nocodazole (Merck); Aurora kinase inhibitor VX-680 (also known as MK-0457, Tozasertib) (Merck); [γ-^32^P] ATP (PerkinElmer, Waltham, MA, USA); Trypsin Gold (Promega, Madison, WI, USA); Glutathione Sepharose 4B beads (GE Healthcare, Chicago, IL, USA); and reduced glutathione (Fujifilm). Recombinant active DAPK3 protein was obtained from Merck.

For nocodazole-arrest and release, cells were treated with 40 ng/ml nocodazole for 14 h, then gently washed twice with prewarmed PBS and cultured for 1 h with the normal culture medium. VX-680 was added to the culture medium, whose final concentration was adjusted to 200 nM.

### General molecular biological techniques

Total RNA extraction and complementary DNA (cDNA) synthesis were performed using the NucleoSpin RNA Kit (Macherey-Nagel, Düren, Germany) and High-Capacity cDNA Reverse Transcription Kit (Thermo Fisher Scientific), respectively. PCR was performed using KAPA HiFi HotStart ReadyMix (Roche, Basel, Switzerland) and a MiniAmpPlus thermal cycler (Thermo Fisher Scientific). For Sanger sequencing, the samples were prepared using the BigDye Terminator v3.1 Cycle Sequencing Kit (Thermo Fisher Scientific) and electrophoresed on a 3500 genetic analyzer (Thermo Fisher Scientific).

### Plasmids

Human full-length LUZP1, AURKB, DAPK3, MYL9, INCENP, BIRC5, and CDCA8 cDNA was amplified via PCR from the HeLa cDNA library and ligated into the pQCXIP vector (Takara Bio, Shiga, Japan) with N-terminal GFP, DsRed, or Flag tag or the pGEX5X-1 vector (Cytiva) with an N-terminal GST tag. The constructs of the LUZP1 deletions or point mutations were generated by partial amplification or site-directed mutagenesis by PCR, respectively.

### Transfection

For plasmid transfection, HeLa and 293T cells were seeded into 35 mm dishes at a density of 2.0 × 10^5^ cells and 8.0 × 10^5^ cells, respectively. On the next day, the cells were transfected with 2 μg plasmid using Lipofectamine 3000 (Thermo Fisher Scientific) for HeLa cells or Polyethylenimine (PEI) Max (Polysciences, Warrington, PA, USA) for 293T cells, according to the manufacturers’ instructions.

For siRNA transfection, 6 × 10^4^ detached HeLa cells were transfected with 20 nM siRNA using Lipofectamine RNAiMAX (Thermo Fisher Scientific) according to the manufacturer’s protocol, then seeded into 35 mm dishes. LUZP1 siRNA #1 (#s15355) and #2 (#s15353) were obtained from Thermo Fisher Scientific. The sequences of the other siRNAs were as follows: KIF20A siRNA: 5’-GGAAAUGUAUGAAGAAAAATT-3’; DAPK3 siRNA: 5’-CUCAGCCGUGAACUACGACTT-3’; and luciferase siRNA: 5’-CUUACGCUGAGUACUUCGATT-3’.

### Immunostaining

Cells were grown on glass coverslips coated with fibronectin (Fujifilm), then fixed with ice cold methanol/acetone (1:1) for 10 min and blocked with PBS that contained 7 % FBS for 30 min. The cells were incubated with the primary antibodies for 1 h, then washed with PBS and incubated with Alexa Fluor 488- and/or 594-labeled secondary antibodies (Thermo Fisher Scientific) for 1 h before being washed with PBS once more. Immunostained images were acquired using a BX60 fluorescence microscope (Olympus, Tokyo, Japan), a BZ-9000 fluorescence microscope (Keyence, Osaka, Japan), an FV1000 laser scanning confocal microscope (Olympus), or an FV3000 laser scanning confocal microscope (Olympus).

### Immunoprecipitation

The cells were lysed with a TNE buffer (25 mM Tris-HCl pH 7.4, 150 mM NaCl, 0.1 % NP-40) or a RIPA buffer (50 mM Tris-HCl, pH 7.4, 150 mM NaCl, 0.1 % SDS, 0.5 % DOC, 1 % NP-40) containing with the cOmplete protease inhibitor cocktail (Roche) and PhosSTOP (Roche), then were centrifuged at 15,000 rpm for 30 min to clear cell debris. The supernatants were incubated with primary antibodies at 4 ºC overnight and then were incubated with protein G-agarose beads (Thermo Fisher Scientific) at 4ºC for 1 h. The beads were collected by centrifugation at 3,000 rpm for 1 min and were washed with each lysis buffer.

### Immunoblotting

Cells were lysed with Laemmli sample buffer and boiled for 5 min. The lysate protein concentrations were determined using the RC-DC Protein Assay (Bio-Rad Laboratories, Hercules, CA, USA). Equal quantities of protein were separated on SDS-PAGE gels and transferred to the PVDF membranes (Merck), which were then blocked with 1 % nonfat skim milk in TBS-T for 30 min, incubated with primary antibodies for 1 h, washed with TBS-T, incubated with HRP-conjugated secondary antibodies, and finally washed with TBS-T. The proteins were visualized by Chemi-Lumi One L (Nacalai Tesque, Kyoto, Japan) and were detected by an Amersham Imager 600 (Cytiva). Band intensities were quantified using ImageJ v1.53p (National Institutes of Health, Bethesda, MD, USA).

### Live cell imaging and Timelapse analysis

HeLa cells were cultured on 35 mm glass base dishes (IWAKI, Japan, Shizuoka) coated with fibronectin and were transfected with GFP and/or DsRed expression plasmid(s). The fluorescence images were acquired 24 h later using an FV1000 or an FV3000 laser scanning confocal microscope (Olympus).

For timelapse analysis, the fluorescence image was taken only at an initial point to determine GFP expressing cells, then the bright-field images were taken for 3 sec intervals for 90 min using an FN1-CSU laser confocal microscope with the incubator at Nagoya University Graduate School of Medicine (Nikon, Tokyo, Japan). Images were analyzed using the MetaMorph Imaging System Software (Universal Imaging, Bedford Hills, NY, USA).

### *In vitro* Kinase assay

The proteins of GST-MYL9 or GST tag alone was purified from *E. coli* using a RIPA buffer, then bound to Glutathione Sepharose 4B beads, washed with a RIPA buffer and then with a kinase buffer (50 mM Tris-HCl pH7.4, 5 mM MgCl_2_, 5 mM MnCl_2_, 5 mM dithiothreitol, 0.01 % Triton X-100) and eluted using 20 mM of reduced glutathione. Flag-LUZP1 FL, Δcc, or Flag tag was transiently expressed into 293T cells, lysed with a RIPA buffer, bound to the bead-conjugated anti-Flag antibodies, and washed with a RIPA buffer and then with a kinase buffer. The RC-DC Protein Assay was used to determine the concentration of all purified proteins. Next 0, 10, 30, 100 ng Flag-LUZP1 FL or Δcc proteins bound to the beads were placed in each tube. At the time, in order to match the bead-to-liquid volume ratio among samples, the purified beads attached with the Flag tag were added and each sample volume was adjusted to 10 μL (beads-to-liquid = 1:1). Then, 20 μL a radio isotope mixture (1 μg of eluted GST-MYL9 or GST, 10 ng of active DAPK3, and 1.0 μCi (0.33 pmol) of [γ^32^-P] ATP) were added into the tube containing with purified LUZP1 and incubated for 30 min at 30°C (final vol., 30 μL). The reaction was terminated by adding the Laemmli sample buffer, then the samples were separated by SDS-PAGE. Finally, the autoradiography image was detected using BAS-5000 (GE Healthcare) and Image Reader BAS-5000 Version 1.8 (Fujifilm), and Multi Gauge Version 3.1 (Fujifilm) software. Band intensities were quantified by using ImageJ v1.53p.

### Protein identification by mass spectrometry

The nocodazole-arrested and released cells or the nocodazole-untreated cells were lysed with TNE buffer, then immunoprecipitated with anti-Flag antibodies. The immunoprecipitated proteins were eluted by boiling for 5 min with 1 % SDS-containing TNE buffer, then the SDS was removed using Detergent Removal Spin Column (Thermo Fisher Scientific). The proteins were reduced, alkylated, and digested using Trypsin Gold (Promega). The peptides were identified at Nagoya University Graduate School of Medicine using the LC-MS/MS system (Paradigm MS4, Michrom Bioresources, Sacramento, CA; HTS-PAL, CTC Analytics AG, Zwingen, Swiss; LTQ Orbitrap XL, Thermo Scientific), and the proteins were identified using the Mascot software package (Matrix Science, London, UK).

### Quantification and Statistical analysis

All statistical analysis were performed using Microsoft Excel (Redmond, WA, USA) and/or using EZR (Saitama Medical Center, Jichi Medical University, Saitama, Japan; Kanda Y, 2013), which is a graphical user interface for R (The R Foundation for Statistical Computing, Vienna, Austria). One-way repeated-measures analysis of variance (ANOVA) with post-hoc Dunnett’s test was used to compare the dose-effect relationships in the kinase assay (Fig. 5D and 5E). Significant differences between the two groups were assessed by performing t-tests. In all statistical analysis, a *p*-value less than 0.05 was considered to be significant.

---

**Supplemental Figure S1.**
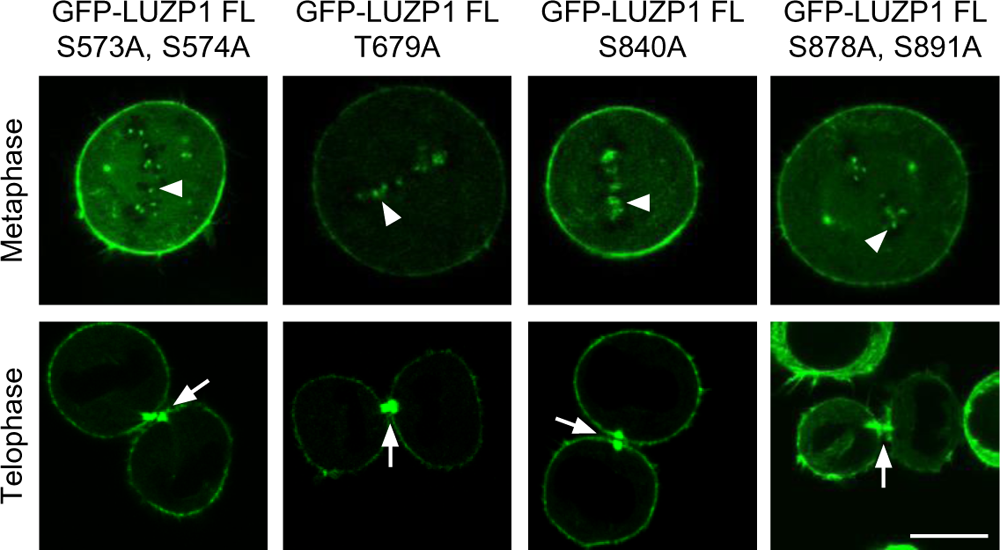
The indicated LUZP1 mutant still localizes at the centromere and the midbody. HeLa cells were transfected with a plasmid encoding indicated GFP-tagged full-length LUZP1 mutant (GFP-LUZP1 FL), then live cell images were taken 24 h later. The arrowheads and the arrows indicate the centromere and the midbody, respectively (scale bar, 10 μm).

**Supplemental Figure S2.**
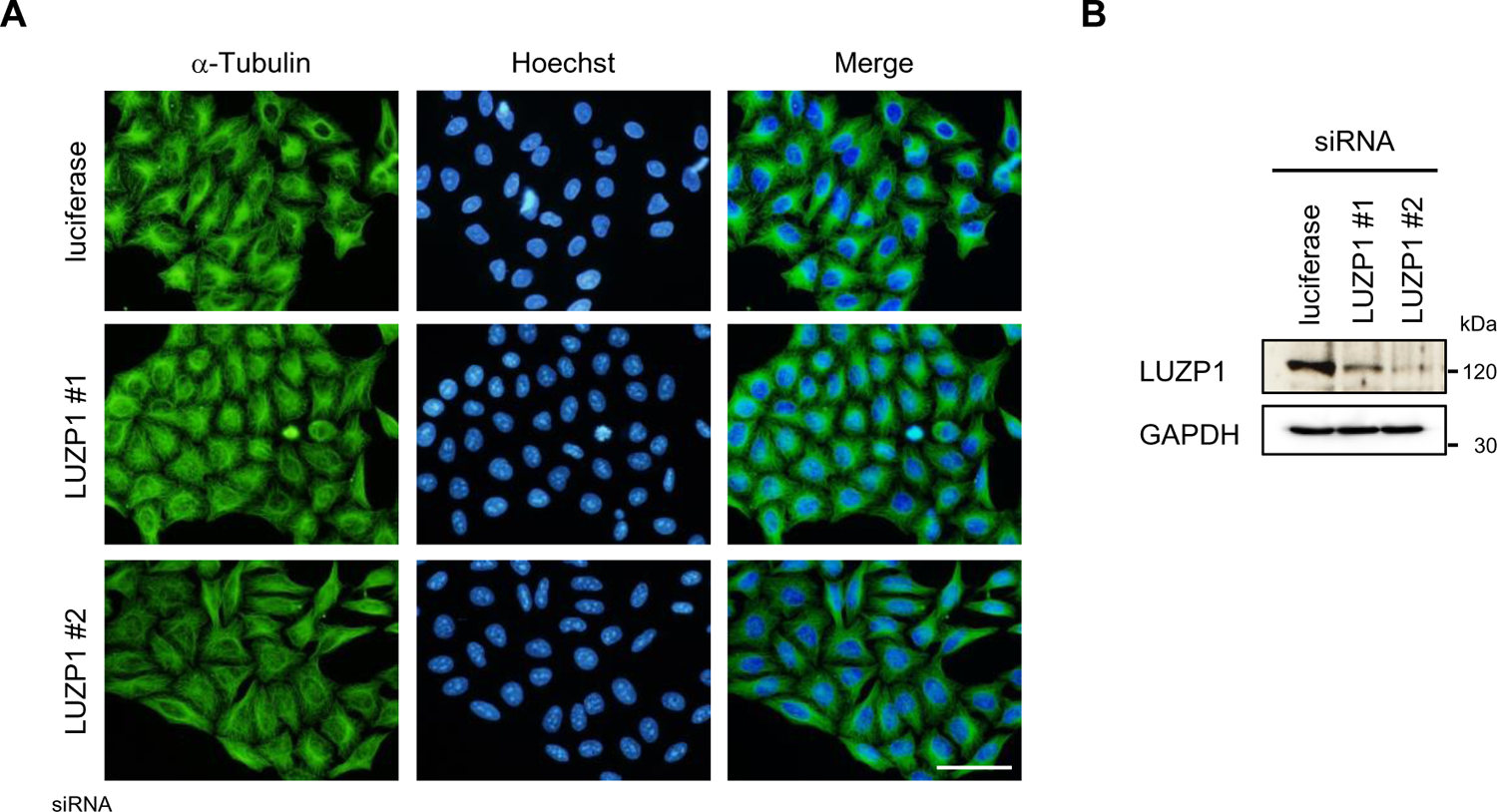
The suppression of LUZP1 does not induce multinucleated cells. (A) HeLa cells were transfected with siRNAs against luciferase or LUZP1, then the cells were immunostained 72 h later with anti-α-Tubulin antibodies and Hoechst (scale bar, 50 μm). (B) HeLa cells were transfected with siRNAs against luciferase or LUZP1 and lysed 72 h later. Next, the cell extracts were immunoblotted with anti-LUZP1 and anti-GAPDH antibodies.

**Supplemental Figure S3.**
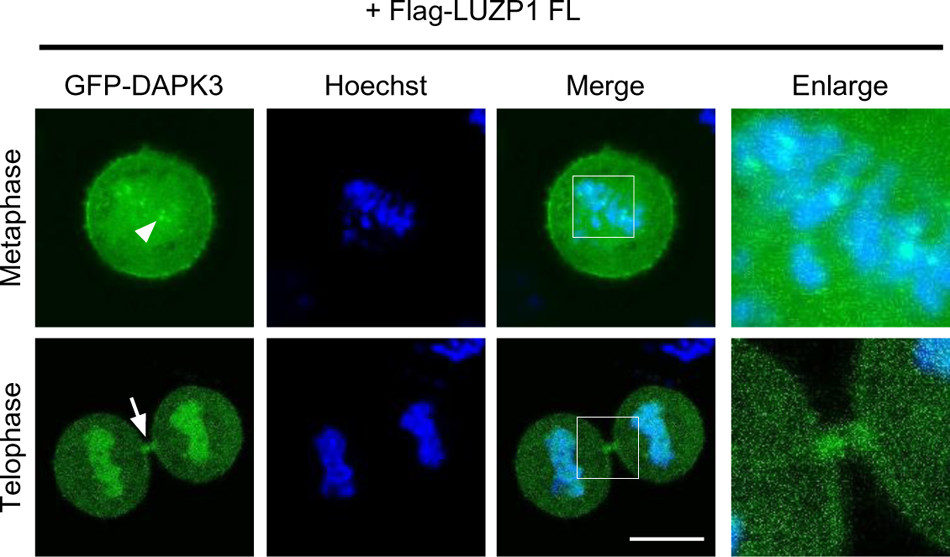
GFP-DAPK3 localized at the centromere in metaphase and at the midbody in telophase when coexpressed Flag-LUZP1 FL. GFP-DAPK3 and Flag-tagged full-length LUZP1 (Flag-LUZP1 FL) were transiently expressed into HeLa cells together, with live images taken 24 h later. The arrowheads and the arrows indicate the centromere and the midbody, respectively. The enlarged image shows the square area (scale bar, 10 μm).

**Supplemental Figure S4.**
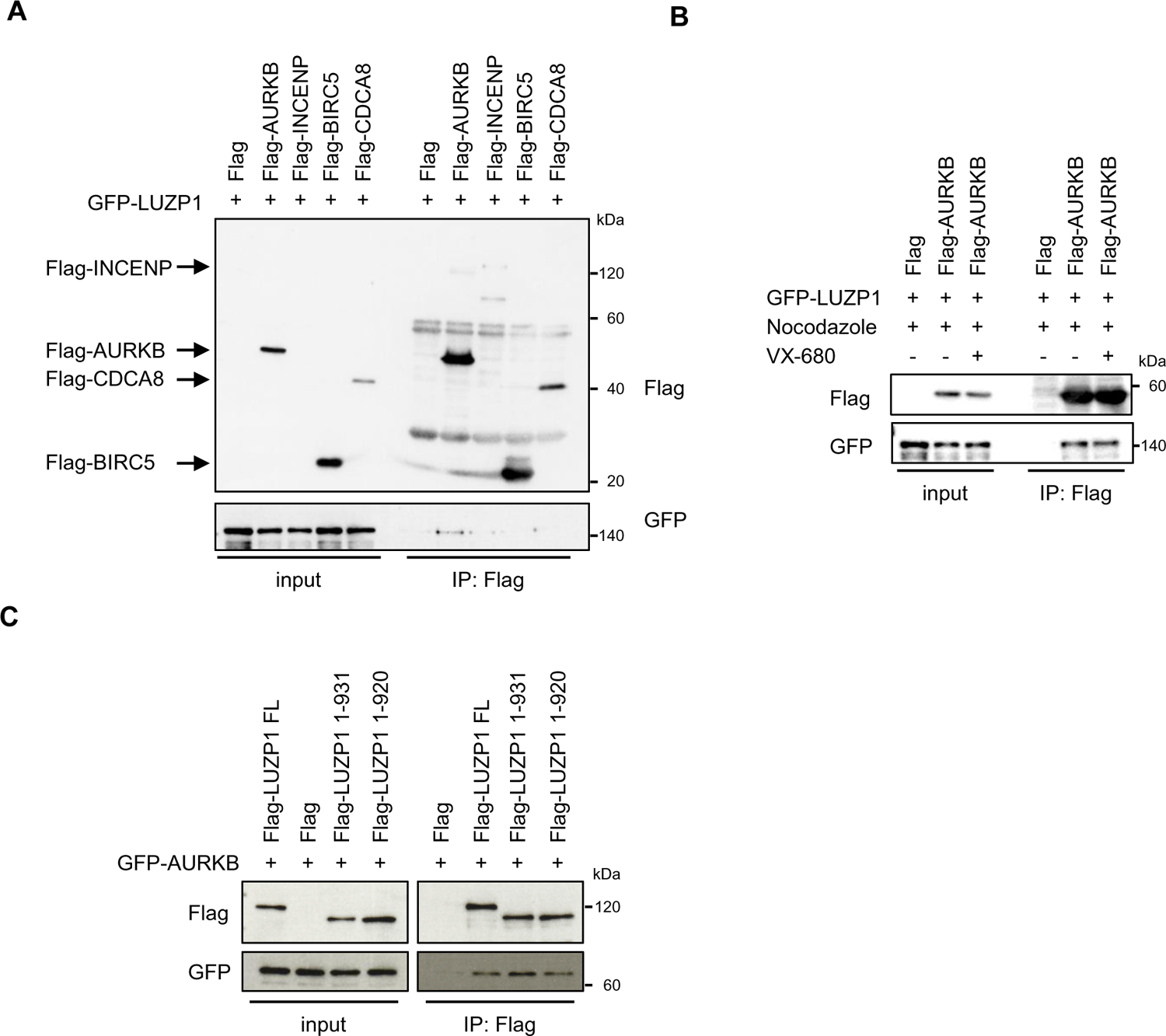
The coimmunoprecipitation analyses between LUZP1 and CPC. (A) GFP-tagged full-length LUZP1 (GFP-LUZP1 FL) and the indicated Flag-tagged full-length CPC component were transiently expressed into 293T cells together and immunoprecipitated with anti-Flag antibodies. The immunoprecipitants were immunoblotted with anti-Flag and anti-GFP antibodies. (B) GFP-LUZP1 FL and Flag-AURKB were transiently expressed into 293T cells together and the cells were treated with nocodazole for 16 h. Next, the cells were additionally treated with or without VX-680 for 1 h and immunoprecipitated with anti-Flag antibodies. The immunoprecipitants were immunoblotted with anti-Flag and anti-GFP antibodies. (C) GFP-AURKB and the indicated Flag-tagged full-length or deletion mutants of LUZP1 were transiently expressed into 293T cells together and immunoprecipitated with anti-Flag antibodies. The immunoprecipitants were immunoblotted with anti-Flag and anti-GFP antibodies.

**Supplemental Table. The raw data of mass spectrometry analysis**.

**Supplemental Movies. Videos of timelapse analysis**.

(S1-S5) The movies correspond to the images in (Fig. 6A). The number shows a stopwatch, which the white and the pink numbers indicate the time in the analysis range and the time before and after following analysis, respectively (scale bar, 10 μm).

