## Supplemental Table for "LUZP1 regulates the constriction velocity of the contractile ring during cytokinesis"

**Supplemental Table. The raw data of mass spectrometry analysis.**

**The nocodazole-untreated cells**

| Swiss prot | | | |
|---|---|---|---|
| Accession | Score | Mass | Description |
| LUZP1_HUMAN | 5745 | 120202 | Leucine zipper protein 1 OS=Homo sapiens GN=LUZP1 PE=1 SV=2 |
| DAPK3_HUMAN | 636 | 52503 | Death-associated protein kinase 3 OS=Homo sapiens GN=DAPK3 PE=1 SV=1 |
| C1QBP_HUMAN | 310 | 31343 | Complement component 1 Q subcomponent-binding protein, mitochondrial OS=Homo sapiens GN=C1QBP PE=1 SV=1 |
| ACTB_HUMAN | 155 | 41710 | Actin, cytoplasmic 1 OS=Homo sapiens GN=ACTB PE=1 SV=1 |
| FILA2_HUMAN | 128 | 247928 | Filaggrin-2 OS=Homo sapiens GN=FLG2 PE=1 SV=1 |
| RHG22_HUMAN | 95 | 76731 | Rho GTPase-activating protein 22 OS=Homo sapiens GN=ARHGAP22 PE=1 |
| ALBU_HUMAN | 95 | 69321 | Serum albumin OS=Homo sapiens GN=ALB PE=1 SV=2 |
| U5S1_HUMAN | 60 | 109366 | 116 kDa U5 small nuclear ribonucleoprotein component OS=Homo sapiens GN=EFTUD2 PE=1 SV=1 |
| HV304_HUMAN | 36 | 12348 | Ig heavy chain V-III region TIL OS=Homo sapiens PE=1 SV=1 |
| SCNND_HUMAN | 35 | 70170 | Amiloride-sensitive sodium channel subunit delta OS=Homo sapiens GN=SCNN1D |
| KS6A6_HUMAN | 35 | 83819 | Ribosomal protein S6 kinase alpha-6 OS=Homo sapiens GN=RPS6KA6 PE=1 |
| RS27A_HUMAN | 34 | 17953 | Ubiquitin-40S ribosomal protein S27a OS=Homo sapiens GN=RPS27A PE=1 |
| CASPC_HUMAN | 32 | 38840 | Inactive caspase-12 OS=Homo sapiens GN=CASP12 PE=2 SV=2 |
| CCD11_HUMAN | 30 | 61796 | Coiled-coil domain-containing protein 11 OS=Homo sapiens GN=CCDC11 PE=1 |
| HSH2D_HUMAN | 30 | 38977 | Hematopoietic SH2 domain-containing protein OS=Homo sapiens GN=HSH2D |
| GPR37_HUMAN | 28 | 67070 | Prosaposin receptor GPR37 OS=Homo sapiens GN=GPR37 PE=1 SV=2 |
| MYLK2_HUMAN | 27 | 64644 | Myosin light chain kinase 2, skeletal/cardiac muscle OS=Homo sapiens GN=MYLK2 PE=1 SV=3 |
| GRP78_HUMAN | 25 | 72288 | 78 kDa glucose-regulated protein OS=Homo sapiens GN=HSPA5 PE=1 SV=2 |
| RIMS1_HUMAN | 24 | 188956 | Regulating synaptic membrane exocytosis protein 1 OS=Homo sapiens GN=RIMS1 |
| NDUB8_HUMAN | 24 | 21751 | NADH dehydrogenase [ubiquinone] 1 beta subcomplex subunit 8, mitochondrial OS=Homo sapiens GN=NDUFB8 PE=1 SV=1 |
| TARB1_HUMAN | 24 | 181559 | Probable methyltransferase TARBP1 OS=Homo sapiens GN=TARBP1 PE=1 SV=1 |
| KDM8_HUMAN | 24 | 47240 | Lysine-specific demethylase 8 OS=Homo sapiens GN=KDM8 PE=1 SV=1 |
| LAMA3_HUMAN | 24 | 366414 | Laminin subunit alpha-3 OS=Homo sapiens GN=LAMA3 PE=1 SV=2 |
| CCD22_HUMAN | 23 | 70712 | Coiled-coil domain-containing protein 22 OS=Homo sapiens GN=CCDC22 PE=1 |
| CR3L4_HUMAN | 23 | 43405 | Cyclic AMP-responsive element-binding protein 3-like protein 4 OS=Homo sapiens GN=CREB3L4 PE=1 SV=1 |
| CC148_HUMAN | 23 | 71031 | Coiled-coil domain-containing protein 148 OS=Homo sapiens GN=CCDC148 |
| FBSP1_HUMAN | 23 | 30613 | F-box/SPRY domain-containing protein 1 OS=Homo sapiens GN=FBXO45 PE=1 |
| ZN503_HUMAN | 22 | 62516 | Zinc finger protein 503 OS=Homo sapiens GN=ZNF503 PE=1 SV=1 |
| GRIK3_HUMAN | 22 | 103970 | Glutamate receptor ionotropic, kainate 3 OS=Homo sapiens GN=GRIK3 PE=2 |
| TIMP4_HUMAN | 22 | 25486 | Metalloproteinase inhibitor 4 OS=Homo sapiens GN=TIMP4 PE=1 SV=1 |
| CCD87_HUMAN | 22 | 96342 | Coiled-coil domain-containing protein 87 OS=Homo sapiens GN=CCDC87 PE=2 |
| SVOP_HUMAN | 22 | 60729 | Synaptic vesicle 2-related protein OS=Homo sapiens GN=SVOP PE=2 SV=1 |
| GDIB_HUMAN | 22 | 50631 | Rab GDP dissociation inhibitor beta OS=Homo sapiens GN=GDI2 PE=1 SV=2 |
| UACA_HUMAN | 22 | 162404 | Uveal autoantigen with coiled-coil domains and ankyrin repeats OS=Homo sapiens GN=UACA PE=1 SV=2 |
| H13_HUMAN | 21 | 22336 | Histone H1.3 OS=Homo sapiens GN=HIST1H1D PE=1 SV=2 |
| LRMP_HUMAN | 21 | 62083 | Lymphoid-restricted membrane protein OS=Homo sapiens GN=LRMP PE=1 SV=3 |
| PKHO2_HUMAN | 21 | 53317 | Pleckstrin homology domain-containing family O member 2 OS=Homo sapiens GN=PLEKHO2 PE=2 SV=1 |
| CAD23_HUMAN | 20 | 369266 | Cadherin-23 OS=Homo sapiens GN=CDH23 PE=1 SV=2 |
| HNRPD_HUMAN | 20 | 38410 | Heterogeneous nuclear ribonucleoprotein D0 OS=Homo sapiens GN=HNRNPD |
| ERO1B_HUMAN | 20 | 53509 | ERO1-like protein beta OS=Homo sapiens GN=ERO1LB PE=1 SV=2 |
| DYH17_HUMAN | 18 | 511460 | Dynein heavy chain 17, axonemal OS=Homo sapiens GN=DNAH17 PE=1 SV=2 |
| LAMA5_HUMAN | 18 | 399479 | Laminin subunit alpha-5 OS=Homo sapiens GN=LAMA5 PE=1 SV=8 |
| DHX8_HUMAN | 17 | 139227 | ATP-dependent RNA helicase DHX8 OS=Homo sapiens GN=DHX8 PE=1 SV=1 |
| HAUS4_HUMAN | 16 | 42373 | HAUS augmin-like complex subunit 4 OS=Homo sapiens GN=HAUS4 PE=1 SV=1 |
| ODFP4_HUMAN | 15 | 29215 | Outer dense fiber protein 4 OS=Homo sapiens GN=ODF4 PE=1 SV=2 |
| COE4_HUMAN | 14 | 64433 | Transcription factor COE4 OS=Homo sapiens GN=EBF4 PE=2 SV=2 |
| ERBB2_HUMAN | 13 | 137821 | Receptor tyrosine-protein kinase erbB-2 OS=Homo sapiens GN=ERBB2 PE=1 |

**The nocodazole-arrested and released cells**

| SwissProt Accession | Score | Mass | Description |
|---|---|---|---|
| LUZP1_HUMAN | 4019 | 120202 | Leucine zipper protein 1 OS=Homo sapiens GN=LUZP1 PE=1 SV=2 |
| ALBU_HUMAN | 602 | 69321 | Serum albumin OS=Homo sapiens GN=ALB PE=1 SV=2 |
| DAPK3_HUMAN | 177 | 52503 | Death-associated protein kinase 3 OS=Homo sapiens GN=DAPK3 PE=1 SV=1 |
| C1QBP_HUMAN | 135 | 31343 | Complement component 1 Q subcomponent-binding protein, mitochondrial OS=Homo sapiens GN=C1QBP PE=1 SV=1 |
| FILA2_HUMAN | 100 | 247928 | Filaggrin-2 OS=Homo sapiens GN=FLG2 PE=1 SV=1 |
| PLAK_HUMAN | 39 | 81693 | Junction plakoglobin OS=Homo sapiens GN=JUP PE=1 SV=3 |
| F214A_HUMAN | 37 | 121594 | Protein FAM214A OS=Homo sapiens GN=FAM214A PE=2 SV=2 |
| SCNND_HUMAN | 36 | 70170 | Amiloride-sensitive sodium channel subunit delta OS=Homo sapiens GN=SCNN1D PE=1 SV=2 |
| LRMP_HUMAN | 33 | 62083 | Lymphoid-restricted membrane protein OS=Homo sapiens GN=LRMP PE=1 SV=3 |
| DSC1_HUMAN | 32 | 99924 | Desmocollin-1 OS=Homo sapiens GN=DSC1 PE=1 SV=2 |
| GDE_HUMAN | 32 | 174652 | Glycogen debranching enzyme OS=Homo sapiens GN=AGL PE=1 SV=3 |
| SPB12_HUMAN | 31 | 46247 | Serpin B12 OS=Homo sapiens GN=SERPINB12 PE=1 SV=1 |
| RHG26_HUMAN | 31 | 92177 | Rho GTPase-activating protein 26 OS=Homo sapiens GN=ARHGAP26 PE=1 |
| MGA_HUMAN | 29 | 209720 | Maltase-glucoamylase, intestinal OS=Homo sapiens GN=MGAM PE=1 SV=5 |
| H13_HUMAN | 28 | 22336 | Histone H1.3 OS=Homo sapiens GN=HIST1H1D PE=1 SV=2 |
| DSG1_HUMAN | 28 | 113676 | Desmoglein-1 OS=Homo sapiens GN=DSG1 PE=1 SV=2 |
| HV304_HUMAN | 27 | 12348 | Ig heavy chain V-III region TIL OS=Homo sapiens PE=1 SV=1 |
| ARI1A_HUMAN | 27 | 241892 | AT-rich interactive domain-containing protein 1A OS=Homo sapiens GN=ARID1A PE=1 SV=3 |
| RT31_HUMAN | 27 | 45290 | 28S ribosomal protein S31, mitochondrial OS=Homo sapiens GN=MRPS31 PE=1 |
| NCOA3_HUMAN | 26 | 155195 | Nuclear receptor coactivator 3 OS=Homo sapiens GN=NCOA3 PE=1 SV=1 |
| KDIS_HUMAN | 25 | 196419 | Kinase D-interacting substrate of 220 kDa OS=Homo sapiens GN=KIDINS220 |
| CASPC_HUMAN | 25 | 38840 | Inactive caspase-12 OS=Homo sapiens GN=CASP12 PE=2 SV=2 |
| RS27A_HUMAN | 25 | 17953 | Ubiquitin-40S ribosomal protein S27a OS=Homo sapiens GN=RPS27A PE=1 |
| HMGX3_HUMA | 24 | 168228 | HMG domain-containing protein 3 OS=Homo sapiens GN=HMGXB3 PE=2 SV=2 |
| ERO1B_HUMAN | 24 | 53509 | ERO1-like protein beta OS=Homo sapiens GN=ERO1LB PE=1 SV=2 |
| S6A16_HUMAN | 24 | 82145 | Orphan sodium- and chloride-dependent neurotransmitter transporter NTT5 OS=Homo sapiens GN=SLC6A16 PE=2 SV=1 |
| KDM8_HUMAN | 24 | 47240 | Lysine-specific demethylase 8 OS=Homo sapiens GN=KDM8 PE=1 SV=1 |
| VIR_HUMAN | 24 | 201898 | Protein virilizer homolog OS=Homo sapiens GN=KIAA1429 PE=1 SV=2 |
| TRIM5_HUMAN | 22 | 56302 | Tripartite motif-containing protein 5 OS=Homo sapiens GN=TRIM5 PE=1 SV=1 |
| CNNM3_HUMA | 22 | 76072 | Metal transporter CNNM3 OS=Homo sapiens GN=CNNM3 PE=1 SV=1 |
| OR4CG_HUMAN | 21 | 34967 | Olfactory receptor 4C16 OS=Homo sapiens GN=OR4C16 PE=2 SV=2 |
| CO6A6_HUMAN | 21 | 247019 | Collagen alpha-6(VI) chain OS=Homo sapiens GN=COL6A6 PE=1 SV=2 |
| LS14B_HUMAN | 21 | 42045 | Protein LSM14 homolog B OS=Homo sapiens GN=LSM14B PE=1 SV=1 |
| RGPA2_HUMAN | 21 | 210636 | Ral GTPase-activating protein subunit alpha-2 OS=Homo sapiens GN=RALGAPA2 PE=1 SV=2 |
| VP13D_HUMAN | 20 | 491606 | Vacuolar protein sorting-associated protein 13D OS=Homo sapiens GN=VPS13D |
| EFC14_HUMAN | 20 | 54997 | EF-hand calcium-binding domain-containing protein 14 OS=Homo sapiens GN=EFCAB14 PE=2 SV=1 |
| MYPC3_HUMAN | 20 | 140674 | Myosin-binding protein C, cardiac-type OS=Homo sapiens GN=MYBPC3 PE=1 |
| CYTA_HUMAN | 20 | 11000 | Cystatin-A OS=Homo sapiens GN=CSTA PE=1 SV=1 |
| S6A15_HUMAN | 19 | 81783 | Sodium-dependent neutral amino acid transporter B(0)AT2 OS=Homo sapiens GN=SLC6A15 PE=1 SV=1 |
| MOCS1_HUMAN | 19 | 70061 | Molybdenum cofactor biosynthesis protein 1 OS=Homo sapiens GN=MOCS1 PE=1 |
| LAMA3_HUMAN | 18 | 366414 | Laminin subunit alpha-3 OS=Homo sapiens GN=LAMA3 PE=1 SV=2 |
| MUC19_HUMAN | 17 | 597790 | Mucin-19 OS=Homo sapiens GN=MUC19 PE=1 SV=2 |
| SDS3_HUMAN | 15 | 38112 | Sin3 histone deacetylase corepressor complex component SDS3 OS=Homo sapiens GN=SUDS3 PE=1 SV=2 |
| XIRP1_HUMAN | 15 | 198439 | Xin actin-binding repeat-containing protein 1 OS=Homo sapiens GN=XIRP1 PE=1 |
| CDSN_HUMAN | 15 | 51490 | Corneodesmosin OS=Homo sapiens GN=CDSN PE=1 SV=3 |
| LIPS_HUMAN | 14 | 116525 | Hormone-sensitive lipase OS=Homo sapiens GN=LIPE PE=1 SV=4 |
| TTC13_HUMAN | 14 | 96751 | Tetratricopeptide repeat protein 13 OS=Homo sapiens GN=TTC13 PE=2 SV=3 |
